# Chronic exposure to TNF reprograms cell signaling pathways in fibroblast-like synoviocytes by establishing long-term inflammatory memory

**DOI:** 10.1101/2020.06.27.171348

**Authors:** Umesh Gangishetti, Sergio Ramirez-Perez, Kyle Jones, Abul Arif, Hicham Drissi, Pallavi Bhattaram

**Affiliations:** Department of Orthopaedics, Emory University School of Medicine, Atlanta, GA 30322, USA; Department of Cell Biology, Emory University School of Medicine, Atlanta, GA 30322, USA; Atlanta VA medical Center, 1670 Clairmont Rd, Decatur, GA 30033

## Abstract

Fibroblast-like synoviocytes (FLS) play a critical role in the pathogenesis of rheumatoid arthritis (RA). Chronic inflammation induces transcriptomic and epigenetic modifications that imparts a persistent catabolic phenotype to the FLS, despite their dissociation from the inflammatory environment. We analyzed high throughput gene expression and chromatin accessibility data from human and mouse FLS from our and other studies available on public repositories, with the goal of identifying the persistently reprogrammed signaling pathways driven by chronic inflammation. We found that the gene expression changes induced by short-term tumor necrosis factor-alpha (TNF) treatment were largely sustained in the FLS exposed to chronic inflammation. These changes that included both activation and repression of gene expression, were accompanied by the remodeling of chromatin accessibility. The sustained activated genes (SAGs) included established pro-inflammatory signaling components known to act at multiple levels of NF-kappaB, STAT and AP-1 signaling cascades. Interestingly, the sustained repressed genes (SRGs) included critical mediators and targets of the BMP signaling pathway. We thus identified sustained repression of BMP signaling as a unique constituent of the long-term inflammatory memory induced by chronic inflammation. We postulate that simultaneous targeting of these activated and repressed signaling pathways may be necessary to combat RA persistence.

## Introduction

Fibroblast-like synoviocytes (FLS) are synovial tissue-resident and specialized mesenchymal cells critical for homeostasis (1). Key features of these cells during homeostasis include the production of extracellular matrix (ECM) components and providing nutrients to the synovial fluid (SF) (1, 2). Healthy synovium is composed of multiple layers of FLS, which forms the synovial lining and sublining through cell-cell contacts (2-4). Inflammatory and pro-resolving mediators are tightly regulated to maintain normal synovium functioning (1). However, in inflammatory and autoimmune diseases such as rheumatoid arthritis (RA), an imbalance between these signals causes homeostasis disruption leading to synovial tissue damage, cartilage destruction and bone degeneration.

In the course of RA presentation, persistent inflammatory signals drive the development of an active inflammatory state, which leads to a persistent aggressive phenotype in FLS. These aggressive FLS are often referred to as transformed FLS (1, 4). The transformed FLS are major source of proinflammatory cytokines, cartilage degrading enzymes as well as the osteoblast activating factor and thereby possess an ability to induce synovitis, arthritis and osteoporosis(1-4). Recent genome wide studies have established that multiple signaling and transcriptional factors with potential pathogenic activities exhibit epigenomic reprogramming in the RA FLS (5-7). The inflammatory milieu in RA synovium promotes an abnormal epigenetic landscape, modulates chromatin accessibility and favors the pathogenic processes observed in RA FLS such as invasion, migration and production of proinflammatory mediators (1, 8, 9). The vital role of tumor necrosis factor-alpha (TNF) in triggering and maintaining an aggressive FLS phenotype has been well documented both in the *in vivo* and *in vitro* studies (10-13). Hence, manipulating TNF-induced sustained chromatin activation in RA FLS represent a treatment opportunity that might aid in combating RA persistence and therapeutic resistance.

Previous studies comparing human RA-FLS to macrophages showed that TNF treatment could induce sustained gene expression in RA-FLS but not in macrophages (10, 12, 14, 15). Recent studies suggest that treatment with TNF prolonged chromatin accessibility and histone modifications for up to 72h in the RA-FLS (10). Other studies have also shown that FLS gain a long-term inflammatory memory due to the exposure to TNF (15). However, the identity of the genes and the pathways that attain a long-term inflammatory memory remain undefined. Furthermore, the duration for which the inflammatory memory persists is also not well understood. In the present study, using an integrative analysis of our own and publicly available epigenomic and transcriptomic data, we determined the identity of the key signaling pathways that are part of the inflammatory memory and likely contribute to the persistent inflammatory phenotype of RA-FLS. We compared the short-term and chronic TNF-induced transcriptomic and epigenetic reprogramming in mouse and human FLS derived from healthy and chronically inflamed arthritic joints. Our results suggest that chronic exposure to TNF contribute to the development of long-term inflammatory memory and thereby switching FLS from passive responders to imprinted aggressors through the activation and repression of critical signaling networks.

## Results

### Overlap between short-term and chronic-TNF mediated transcriptomic changes in mouse FLS

We compiled RNA-sequencing (RNA-seq) data from multiple sources; where normal primary mouse FLS (mFLS) between passages 3-5 were treated with or without recombinant human TNF for 16, 24 and 48 hours from independent data sets generated in our laboratory and public data sets available on NCBI-Sequence Read Archive (**Table S1**). We performed differential gene expression analysis to identify genes that were upregulated or downregulated by 2-fold upon TNF treatment. Our analysis revealed upregulation of 925, 279 and 1519 genes after 16, 24 and 48 hour of TNF treatment respectively. We also identified 1398, 123 and 1807 genes downregulated by 2-fold at each of these time points (**Fig S1**). In order to determine whether the short-term TNF-induced transcriptomic changes overlapped with that of gene expression changes resulting from long-term TNF treatment, we obtained RNA-seq data from primary FLS cultured from 11-week-old wild-type and TNF-transgenic mice (Tg197) (16). The results revealed that 244 genes were upregulated and 81 genes were downregulated by 2-fold in the FLS from Tg197 mice. We then compared the genes differentially regulated by short-term and chronic TNF exposure. We thus identified that 80% (195 of 244) of the genes upregulated in FLS exposed to chronic TNF conditions were also upregulated in at least one of the short-term TNF exposures (**Fig 1A, Table S2**). These genes were designated as sustained activated genes in mouse FLS or mSAGs. Likewise, comparative analysis of the downregulated genes identified 80% (65 of 81) of the genes downregulated in FLS exposed to chronic TNF conditions were also downregulated in at least one of the short-term TNF exposures (**Fig 1B, Table S3**). These genes were designated as sustained repressed genes in mouse FLS or mSRGs. Heatmap analysis further verified the differential expression of mSAGs and mSRGs in all conditions (**Fig 1C and D**). Ingenuity pathway analysis revealed that the top ten disease and functions enriched for the mSAGs were primarily involved in connective tissue, skeletal and immunological diseases. In contrast, the disease and functions associated with the mSRGs were related to tissue development, growth and metabolism (**Fig 1E**). Ingenuity canonical pathway analysis revealed that the molecular mechanisms regulated by mSAGs were involved in inflammatory signaling whereas the mechanisms regulated by mSRGs were predominantly involved in the molecular pathways required for stem cell proliferation and tissue differentiation and BMP signaling in addition to RA and osteoarthritis (OA) pathways (**Fig 1F)**. Together, these data demonstrated that removing FLS from the inflammatory environment in the arthritic joints and placing them under nutrient rich *in vitro* culture conditions does not revert these sustained gene expression changes. This also supports the notion that, chronic inflammation induces a “long-term inflammatory memory” in the FLS transcriptome.

**Figure 1.**
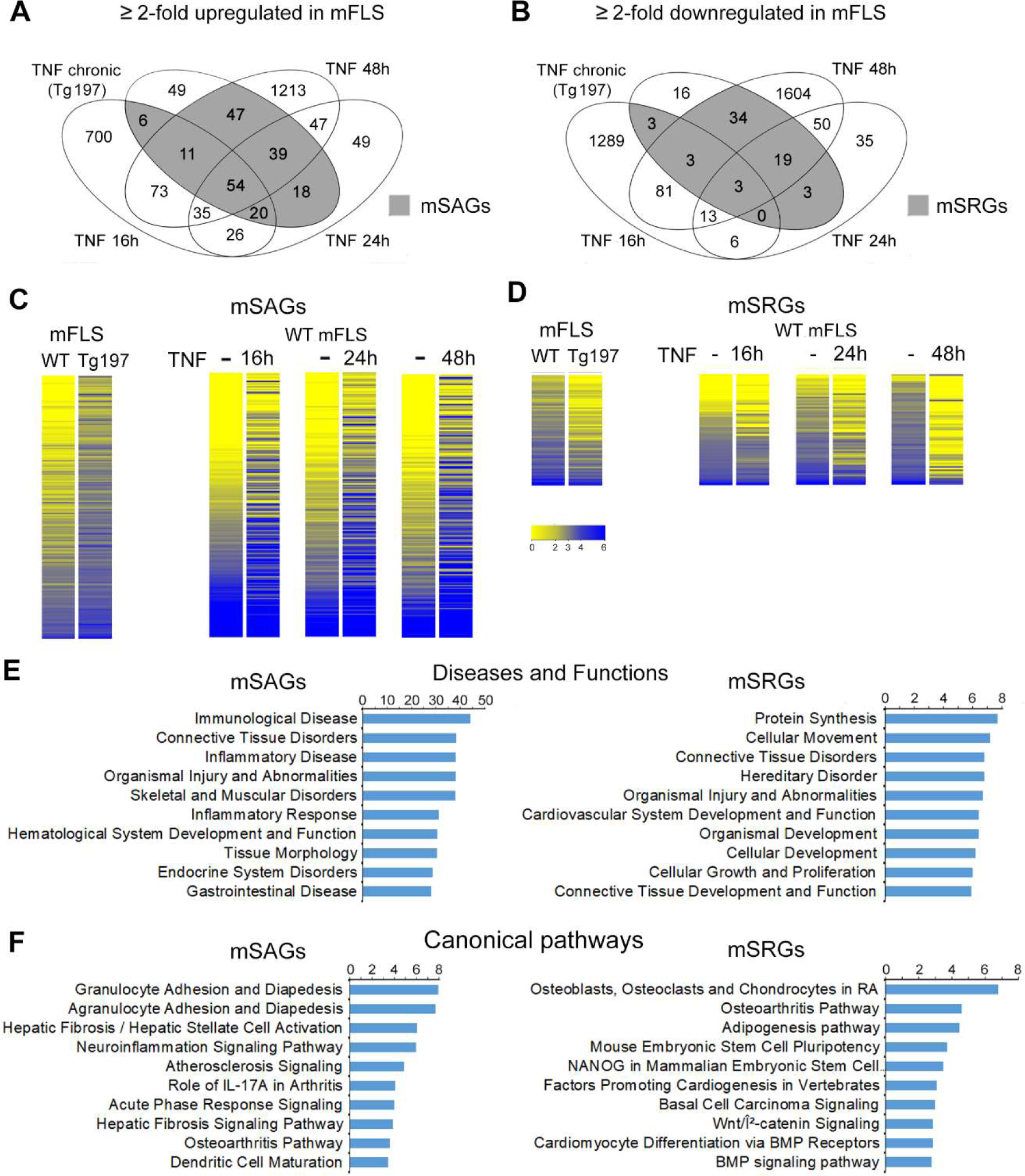
Chronic TNF induces sustained gene expression changes in mouse Fibroblast-like synoviocytes (mFLS). (A) Venn diagram showing the number of upregulated genes by 2-fold in the short-term TNF treatment of wild-type mouse FLS (TNF 10ng/mL; 16h, 24h and 48h) and Tg197 human TNF-expressing transgenic mice (chronic TNF exposure). Grey shaded area represents the overlapping genes in FLS of the short-term and chronic-TNF, and is referred as sustained activated genes in mouse (mSAGs). (B) Venn diagram showing the number of downregulated genes in each condition described in A. Grey shaded area shows the overlapping genes between short-term and chronic-TNF, and is referred as sustained repressed genes in mouse (mSRGs). (C and D) Heatmap of mSAGs and mSRGs. Each line represents one gene and is color-coded based on normalized expression levels. (E) Top 10 diseases and functions regulated by mSAGs and mSRGs identified by IPA analysis. (F) Top 10 canonical pathways regulated by mSAGs and mSRGs identified by IPA analysis.

### Overlap between short-term and chronic-TNF mediated epigenomic changes in mouse FLS

In order to know the extent to which short-term TNF treatment could change the availability of open chromatin in mouse FLS, we performed ATAC-sequencing experiment from mFLS treated without or with TNF for 16h. ATAC-sequencing identified 12,548 peaks in untreated samples while 36,881 peaks were identified in the TNF-treated indicating that TNF enhances chromatin accessibility. The peaks from TNF-treated FLS were associated with transcription start sites of genes at a much higher frequency than in untreated FLS control (**Fig 2A**). In order to study chromatin accessibility changes due to chronic inflammation, we analyzed DNase-seq data from FLS of wild-type control and a mouse model of inflammatory arthritis that were generated as a part of the human and epigenome database ‘DEEP’ http://deep.dkfz.de/ (**Table S1**). Peak calling revealed 27,984 regions to be accessible to DNase I digestion, which as expected increased to 41,665 accessible regions under inflammatory conditions (**Fig 2B**). Moreover, the genes assigned to the peaks identified by ATAC-seq and the DNase-seq overlapped with the gene expression changes of their respective conditions. Overlap was also observed in both genes that were upregulated and downregulated by TNF (**Fig 2C and D**). Together these data suggest that the changes in chromatin accessibility can lead to the recruitment of transcriptional activation complexes, as well as transcriptional repression complexes to the genomic regulatory regions.

**Figure 2.**
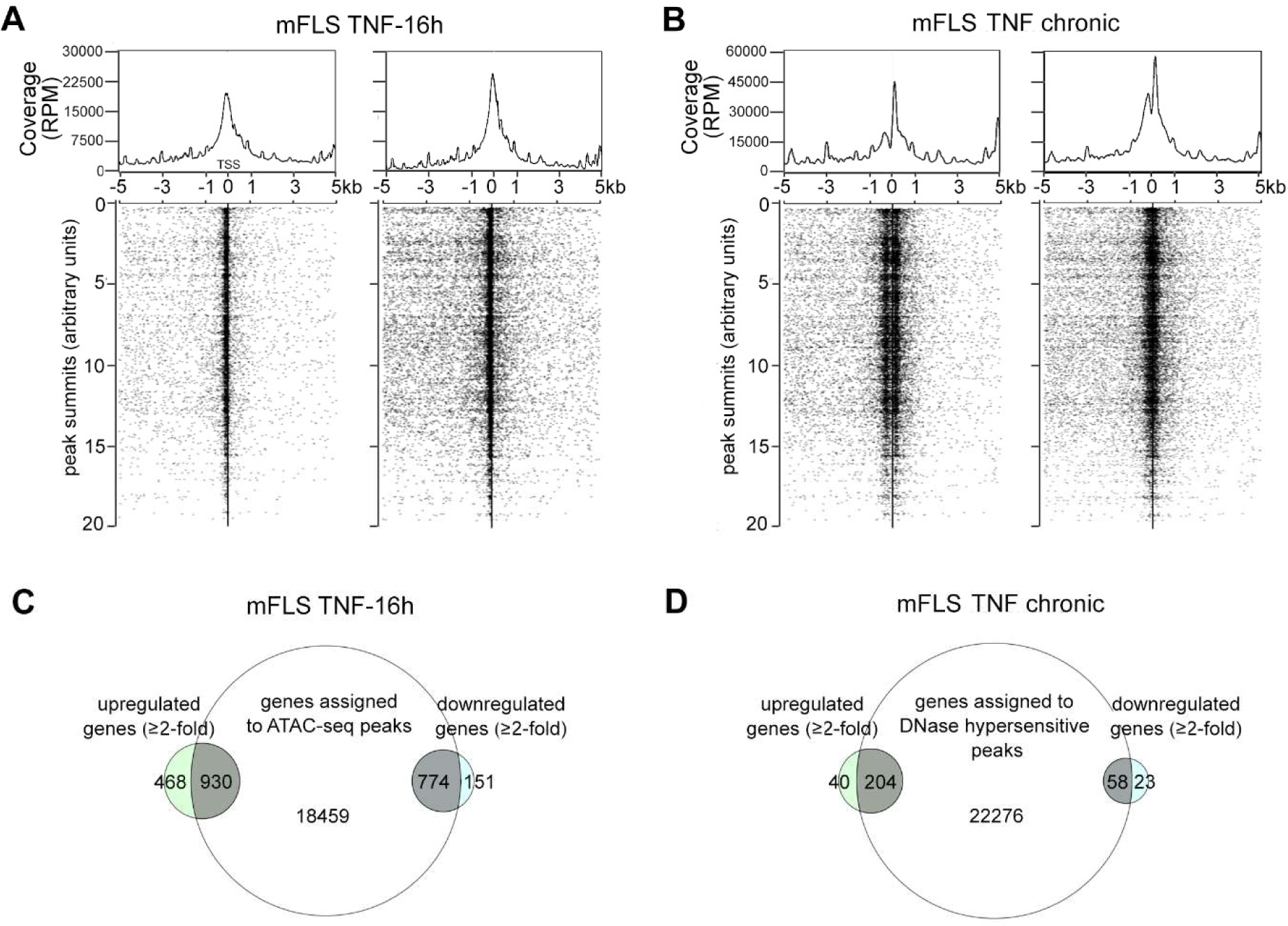
TNF-induced chromatin accessibility changes overlap with transcriptomic changes. **(A)** TSS plot of mFLS treated with or without 10 ng/mL TNF for 16h showing the distribution of ATAC-seq peaks 5 kb upstream and downstream of the transcription start sites (TSS). Each dot represents one peak assigned to a unique gene. **(B)** TSS plot of mFLS isolated 13-week old wild-type control mouse paws and chronically inflamed paws of RA mouse showing the distribution of DNase-seq peaks 5 kb upstream and downstream of the transcription start sites (TSS). Each dot represents one peak assigned to a unique gene. **(C)** Venn diagram showing the overlap between differentially expressed genes and genes assigned to ATAC-seq peaks. **(D)** Venn diagram showing the overlap between differentially expressed genes and genes assigned to DNase-seq.

### Identification of sustained activated transcriptomic and epigenomic reprogramming in mouse and human RA FLS

In order to determine whether the genes exhibiting sustained differential expression between short-term and chronic TNF-exposure also exhibited sustained changes in chromatin accessibility, we compared the genes assigned to the chromatin accessibility peaks with that of the mSAGs (**Fig 1A**). We observed that 72.3% (141 of 195) mSAGs contained accessible chromatin within 5 kb of their TSS in both short-term and chronic-TNF exposure conditions (**Fig 3A**). We next determined whether these 141 mSAGs were also activated in human RA FLS. For this, we analyzed RNA-seq data sets from the SRA database, where patient-derived RA FLS were treated with TNF (**Table S1**). We found that 825 and 946 genes were upregulated by > 2-fold at 8h and 24h of TNF treatment (**Fig S1**). There was a 75% overlap (590 of 787) between genes upregulated at 8h and 24h. These overlapping upregulated genes also contained 52 of the 141 SAGs that were determined to exhibit both epigenetic and transcriptomic reprogramming (**Fig 3B and Table S4**). Heatmap analysis of normalized intensities of the identified human SAGs shows the differential expression between the treated and untreated RA FLS (**Fig 3C**). In order to determine the functional relationship between the hSAGs, we performed STRING network analysis, a computational method used to detect protein-protein interaction networks based on functional enrichment analysis. We found that the hSAGs exhibited strong interactions, with the canonical and non-canonical NF-κB signaling transcription factor RELB, interacting with maximum number of hSAGs (**Fig 3D**). In addition, we found multiple genes associated with pro-inflammatory pathways such as genes encoding for the components of prostaglandin synthesis pathway including prostaglandin E synthase (*PTGES*) and prostaglandin I2 receptor (*PTGIR*), pro-angiogenic vascular endothelial growth factor C (*VEGFC*), a member of the poly(ADP-ribose) polymerase protein family (*PARP14*) as well as the Fas cell surface death receptor (*FAS*). We next performed a prediction analysis to determine the transcription factors that are likely to regulate the 52 SAGs, using the transcription factor enrichment analysis ChEA3 (17). This analysis prioritizes transcription factors based on the overlap between given lists of differentially expressed genes based on the putative targets as determined by ChIP-seq experiments from ENCODE projects. The results revealed RELA, STAT1, STAT2, JUN and JUND among the top 10 transcription factors predicted to regulate the expression of SAGs (**Fig 3E**). Finally, we visualized the RNA-seq and chromatin accessibility sequencing peak profiles at a few representative mSAG (*Nfkbia* and *Bric3*; **Fig 4A**) and hSAGs (*RELB, FAS* and *IRF*; **Fig 4B**). *Nfkbia* (codes for inhibitor kappa alpha protein), *Birc3* (baculoviral IAP repeat containing 3), *RELB (*encodes for p50 NF-kB transcription factor*), FAS* (encodes for TNF superfamily cell surface receptor involved in programmed cell death) and *IRF1* (interferon regulatory factor 1, encodes for a transcriptional regulator of immune responses) are examples of components of various proinflammatory signaling cascades. The RNA-seq peak intensities at the exons of each of these genes was higher in both short-term and chronic TNF conditions, compared to the untreated controls. Concomitantly, the intensity of ATAC-seq and DNase-seq peaks at the promoters of these genes was also increased by both short-term and chronic TNF-exposure.

**Figure 3.**
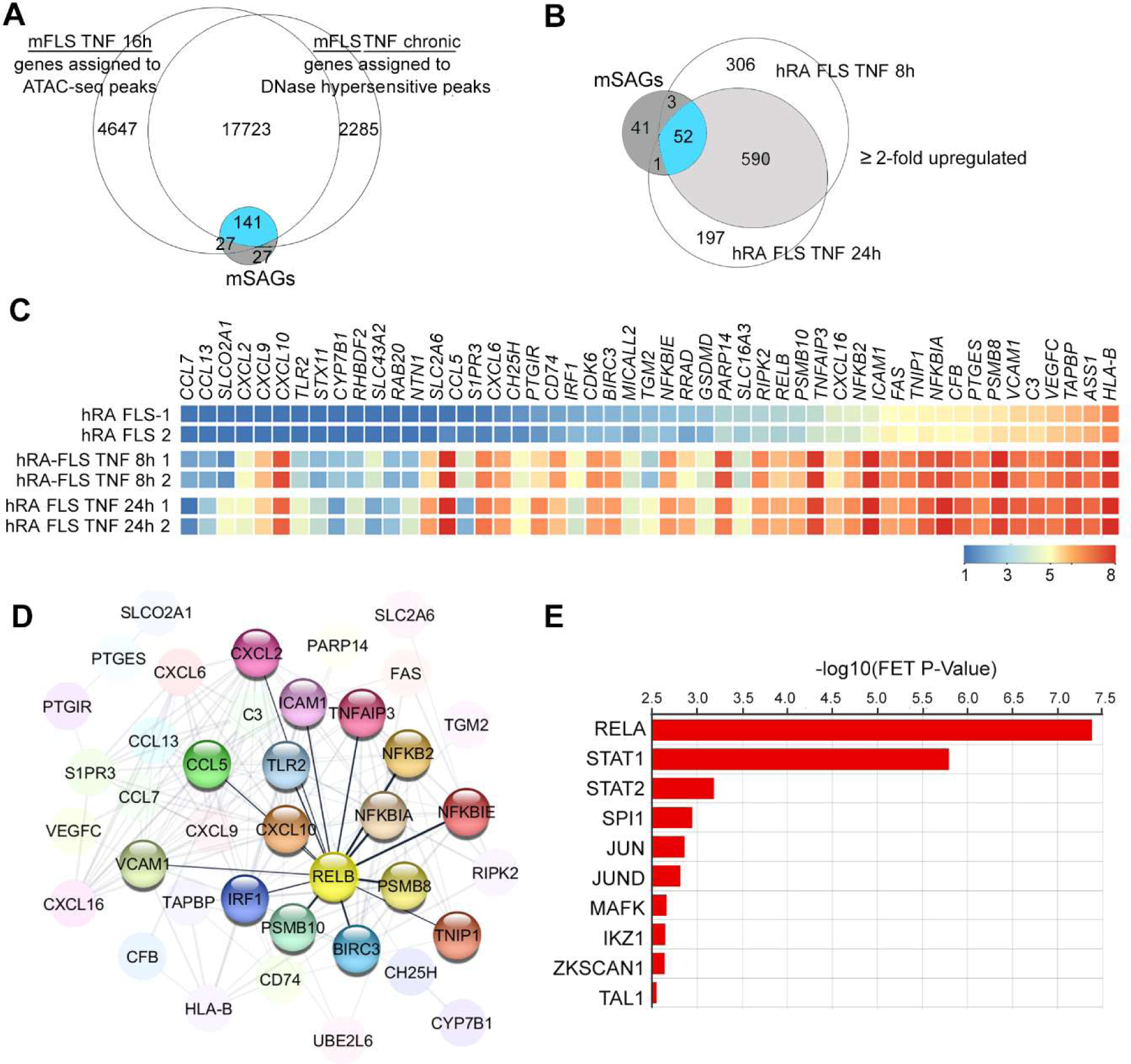
SAGs regulate pro-inflammatory signaling networks in mouse and human FLS. **(A)** Venn diagram showing the overlap between genes assigned to ATAC-seq and DNase-seq peaks in short-term (16h) and chronic-TNF conditions with that of mSAGs identified from Fig 1A. Overlapping SAGs are shown in blue. **(B)** Venn-diagram showing the overlap between gene upregulated by > 2-fold in human RA-FLS treated with 10 ng/mL recombinant TNF for 8h and 24h with that of 141 mSAGs identified in Fig 3A. 52 overlapping human SAGs are shown in blue. **(C)** Heatmap of hSAGs. Each box represents one gene and is color-coded based on normalized expression levels. **(D)** Functional protein-protein interaction between hSAGs by STRING analysis. Proteins interacted with RELB are highlighted. **(E)** Top 10 transcriptional factors predicted to regulate hSAGs expression identified by ChEA3 online tool.

**Figure 4.**
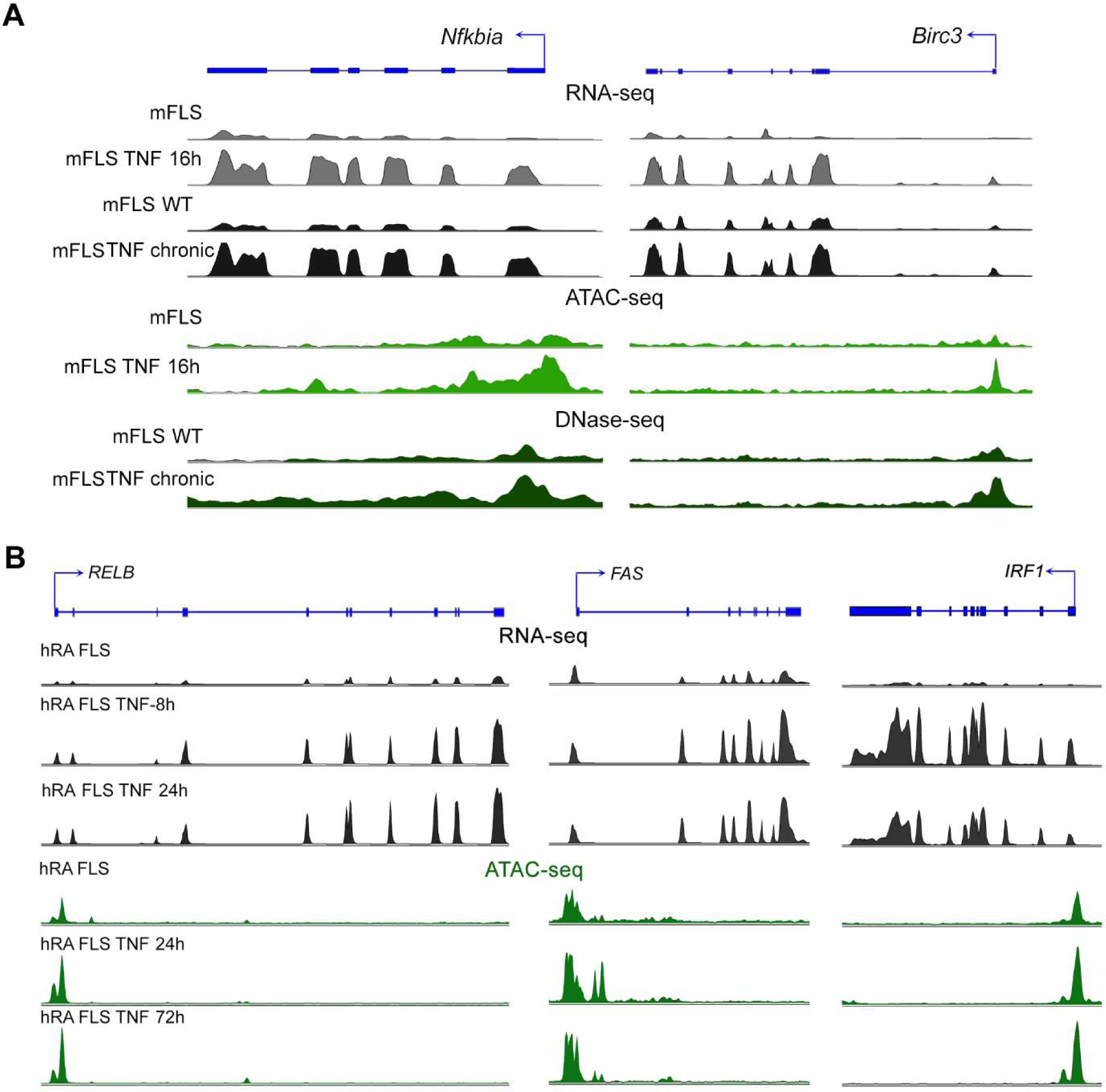
Transcriptomic and chromatin accessibility profile changes at mouse and human SAGs. **(A)** RNA-seq, ATAC-seq and DNase-seq peak profiles in mFLS for the indicated conditions at *Nfkbia* and *Birc3* genomic loci. **(B)** RNA-seq and ATAC-seq peak profiles in hRA-FLS for the indicated conditions at *RELB, FAS* and *IRF1* genomic loci.

### Identification of sustained repressed transcriptomic and epigenomic reprogramming in human RA-FLS

Our next goal was to identify whether the repressed mSRGs were associated with sustained changes in chromatin accessibility. We observed that 52.3% (34 of 65) mSRGs contained accessible chromatin within 5 kb of their TSS in both short-term and chronic-TNF exposure conditions (**Fig 5A**). We then wanted to ascertain if the mSRGs were also repressed in human RA FLS. We, therefore, analyzed RNA-seq data sets in which human RA-FLS were treated with TNF. The results revealed that 1261 and 854 genes were downregulated by > 2-fold after 8h and 24h of TNF treatment respectively (**Fig S1**) and found that 65% (577 of 851) of the genes downregulated at 8h of TNF treatment remained downregulated even at 24h TNF treatment. Of these 34 mSRGs that were repressed at the epigenomic and transcriptomic level by chronic inflammation in mice, only 7 genes (20%) were confirmed to be human SRGs as they were downregulated by > 2-fold in human RA FLS treated with TNF for 8h and 24h (**Fig 5B and Table S4**). The differential expression of these seven hSRGs in RA FLS treated with TNF are also shown in the heat map (**Fig 5C**). Functional protein-protein analysis by STRING pathways, identified that the proteins encoded by 3 of the 7 hSRGs exhibited unexpected functional interactions with BMP signaling (**Fig 5D)**. Bone morphogenic protein ligand (*BMP4*), its downstream transcriptional effector *SMAD9* (also known as SMAD8) and the BMP signaling target and transcriptional regulator from the family of the inhibitor of DNA binding and cell differentiation proteins (*ID4)* were a part of this predicted network. CheA3 transcription factor enrichment analysis tool predicted that the transcriptional repressor CCCTC-Binding Factor (CTCF) (18) is one of the top candidates likely to downregulate the expression of hSRGs in response to chronic inflammation (**Fig 5E**). Finally, we visualized the RNA-seq and chromatin accessibility sequencing peak profiles at some of the representative mSRGs (**Fig 6A**) and hSRGs (**Fig 6B**). TNF-exposure resulted in a decrease in the intensity of RNA-seq peaks at the exons of these genes. The intensity of ATAC-seq and DNase-seq peaks at the promoter regions mSRGs were also increased by TNF treatment. These data support the notion that TNF recruits transcriptional repressors such as CTCF to the regulatory regions of SRGs to repress their expression. However, the intensity of the ATAC-seq peaks at the promoters of the hSRGs in RA FLS was only modestly increased by TNF treatment, in spite of a reduction in the RNA-seq peaks. These data suggest that inflammatory memory of the chromatin landscape at SRGs genomic loci was already established in the human RA FLS and additional TNF treatment was almost inconsequential.

**Figure 5.**
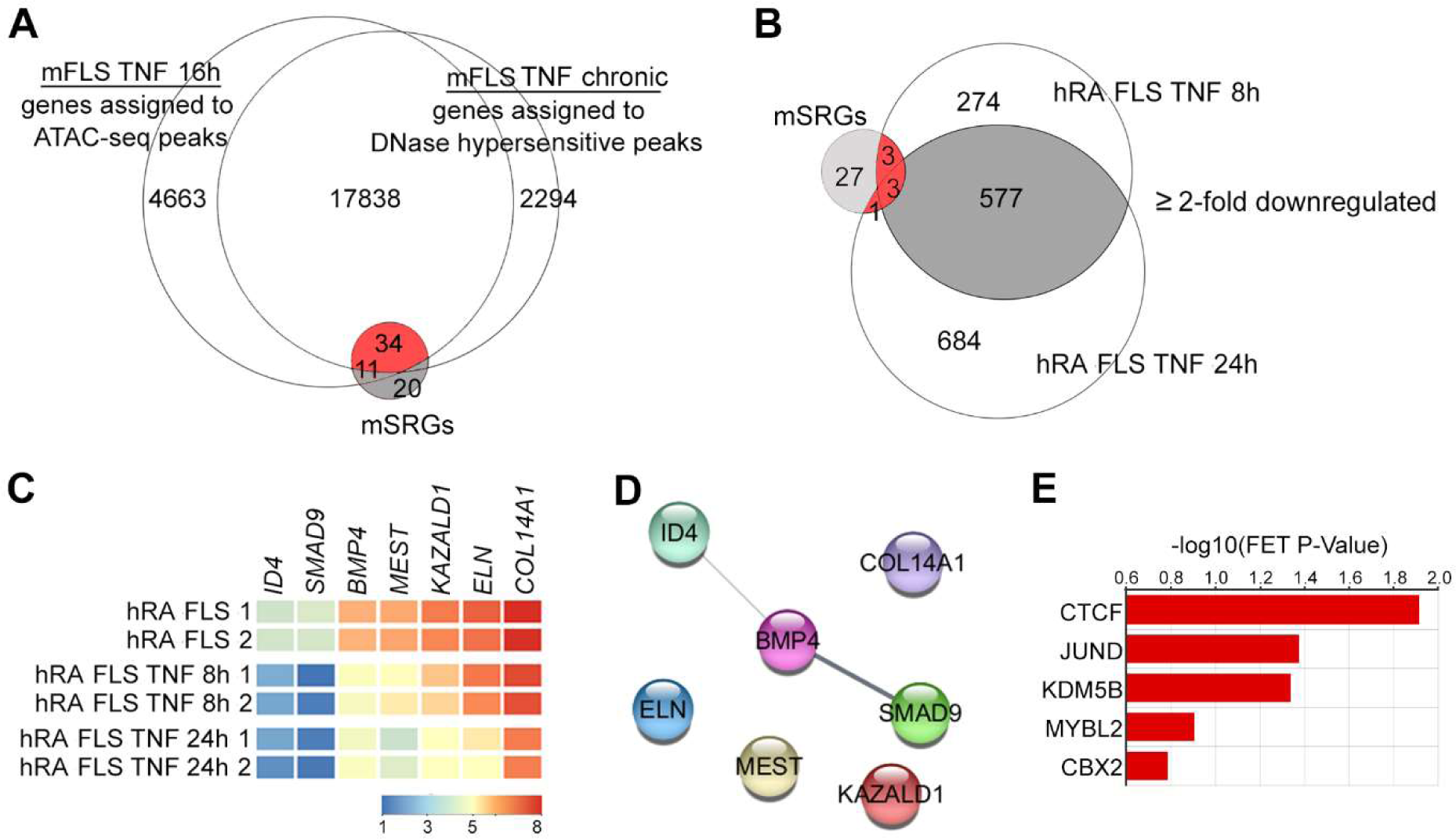
SRGs regulate BMP signaling in mouse and human FLS. **(A)** Venn diagram showing the overlap between genes assigned to ATAC-seq and DNase-seq peaks in short-term (16h) and chronic-TNF conditions with that of mSRGs identified from Fig 1B. Overlapping SRGs are shown in red. **(B)** Venn-diagram showing the overlap between genes downregulated by > 2-fold in human RA-FLS treated with 10ng/mL recombinant TNF for 8h and 24h with that of 34 mSAGs identified in Fig 5A. 7 overlapping human SRGs are shown in red. **(C)** Heatmap of hSRGs. Each box represents one gene and is color-coded based on normalized expression levels. **(D)** Functional protein-protein interaction between hSRGs by STRING analysis. **(E)** Top 5 transcriptional factors predicted to regulate hSAGs expression identified by ChEA3 online tool.

**Figure 6.**
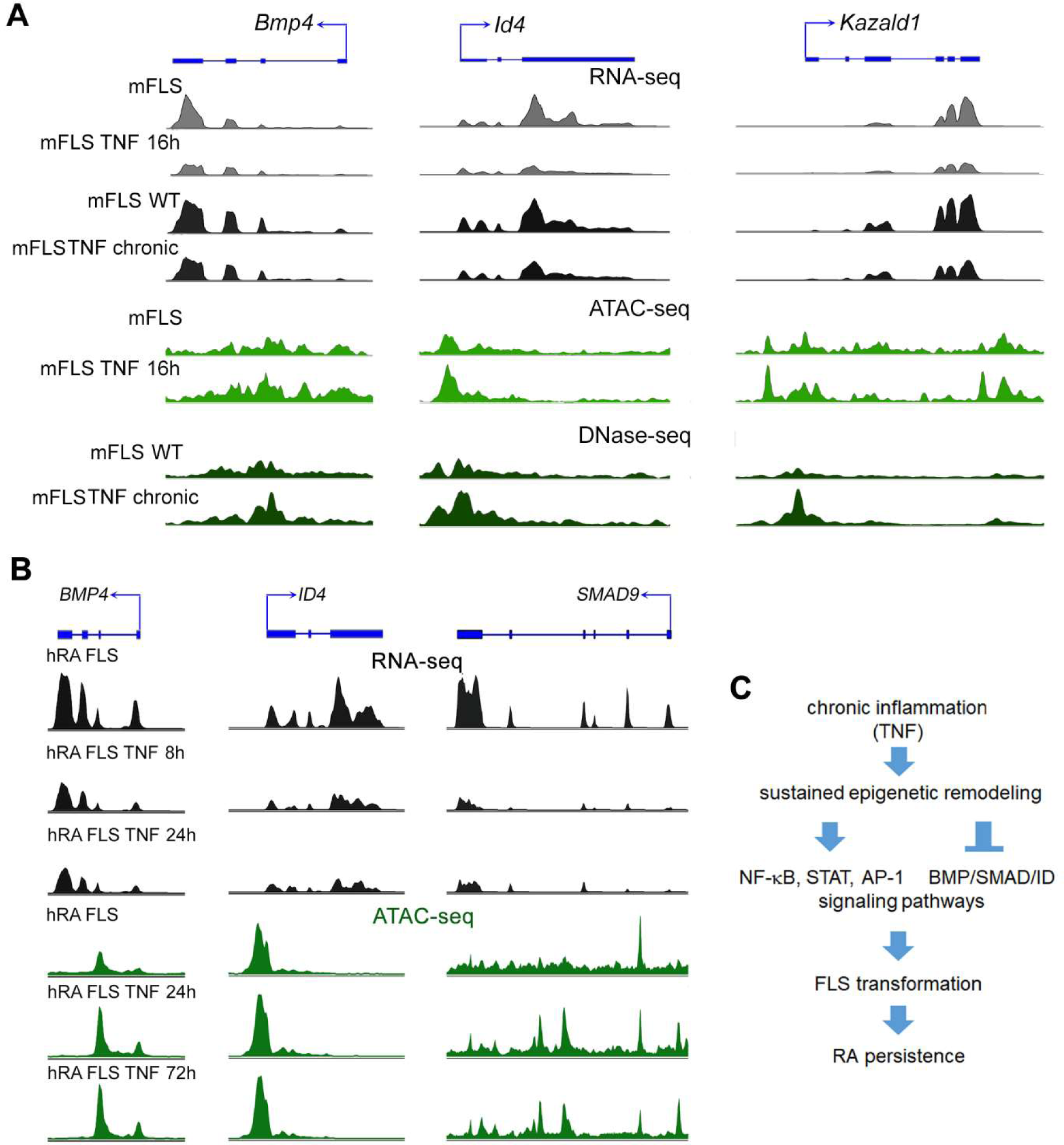
Transcriptomic and chromatin accessibility profile changes at mouse and human SAGs. **(A)** RNA-seq, ATAC-seq and DNase-seq peak profiles in mFLS for the indicated conditions at *Bmp4, Id4* and *Kazald1* genomic loci. **(B)** RNA-seq and ATAC-seq peak profiles in hRA-FLS for the indicated conditions at *BMP4, ID4* and *SMAD9* genomic loci. **(C)** Schematic showing persistent and long-term reprogramming of signaling pathways in FLS exposed to chronic TNF exposure.

## Discussion

Epigenetic regulation of the genome is one of the primary means by which the genetic code is altered to control cellular functions. Mechanisms such as histone modifications, DNA methylation and chromatin remodeling together govern the epigenetic landscape that dictate the outcome of cell fate (19). Epigenetics reprogramming of FLS in particular is suggested to be a major mediator of FLS transformation, RA persistence and therapeutic resistance (5, 8). Recent large-scale genome-wide studies have incontrovertibly established that FLS transformation is associated with epigenetic-remodeling. However, which of these changes are reversible and which of them are permanent is not fully understood (5, 14, 16) A recent study identified that multiple pro-inflammatory genes in human RA FLS escaped transcriptional repression for at least up to period of 72h of TNF treatment. These escaped genes were suggested to contribute to the persistence of synovitis observed in RA patients. This study also reported that the reprogramming seems to be restricted to RA FLS and was not observed in macrophages due to their tolerogenic mechanisms, which suppress the expression of TNF-inducible genes (14). We aimed to determine whether the absence of tolerogenic mechanisms persists for a long-term and if escape from transcriptional repression can also be observed in FLS from TNF-transgenic mouse model of inflammatory arthritis. The irreversible transformation of FLS behavior directly depends on the signaling cascades and transcriptional networks that are reprogrammed due to the epigenetic mechanisms (5). We set out to identify the signaling and transcriptional networks that are specifically susceptible to long-term modulation. It was previously reported that the genomic responses of FLS from TNF-transgenic mice greatly resemble those of RA FLS from humans, suggesting that molecular mechanisms underlying long-term inflammatory memory overlap between mouse and humans (16). We therefore performed a comparative analysis of next-generation sequencing data from 28 different mFLS samples that were subjected to TNF either for short periods or for chronic exposure in the inflamed joints. We further compared these results with the genome-wide data from 9 human RA FLS samples treated with TNF for varying time periods. Some of these data were generated in our laboratory and others were accessed from the public repositories (14, 16). This exercise provided a critical insight into the molecular understanding of the sustained epigenetic remodeling. Importantly, it also led to the identification of the signaling cascades both activated and repressed by these epigenomic changes (**Fig 6C**). Our pathway and functional prediction analysis for mSAGs also revealed their strong association with inflammatory diseases and pathways, which aligns with the established roles for pro-inflammatory signaling and transcriptional networks in autoimmune joint diseases. Hence, our data showing that NF-κB, STAT and AP-1 signaling pathways exhibit sustained activation was expected (2, 4, 20). However, to which extent the individual components of these signaling pathways exhibit sustained activation has not been previously recognized. We observed that the RA FLS accumulate a large number of epigenetic modifications to ensure persistent and increased activation of the NF-κB signaling pathway. For instance, the sustained activation of *RELB* would ensure that both canonical and non-canonical arms of the NF-κB pathway would show an enhanced response to inflammatory ligands in the RA FLS (21). The enhanced activation of the NF-κB pathway is likely further increased by the sustained activation of receptors, such as toll-like receptor (TLR2) and intermediate components such as NF-κB subunit 2 (NFKB2, also called p100), NF-κB inhibitor alpha (NFKBIA, also called IκB-Alpha) and NF-κB inhibitor epsilon (NFKBIE, also called IκB-epsilon). Other SAGs that directly and indirectly contribute to the sustained and exacerbated NF-κB signaling included the TNFAIP3, which encodes the deubiquitinating enzyme A20, that plays a role in the NF-κB non-canonical activation (22). The C-C motif and multiple C-X-C motif chemokine ligands (23) and prostaglandin synthesis pathway components were other additional pro-inflammatory pathways that show sustained activation in the RA FLS (24). Together these data have uncovered that FLS attain a robust long-term inflammatory memory in response to chronic inflammation. The SAGs identified to possess “long-term inflammatory memory” are likely to govern the aggravated and sustained responses to inflammatory stimuli in the transformed RA-FLS.

Notably, we observed that chronic TNF induces certain genes to undergo sustained repression, which we refer to as SRGs. At the functional level, the SRGs were predicted to function in pathways regulating stem cell proliferation, growth and differentiation. Interestingly, our study also suggests that SRGs may regulate BMP signaling pathway. In fact, BMP signaling was found to be active in several rheumatic diseases such as oligoarticular juvenile idiopathic arthritis (JIA) and RA. However, the expression of key molecules involved in this signaling pathway seems to differ with disease context and its implications in RA are not completely understood (25, 26). A previous study reported that BMP4 is downregulated in the synovial tissue of RA patients (27). Others reported that BMP4 plays a critical role in upregulating the expression of SMAD9 due to multiple BMP-responsive elements located at the promoter region of *SMAD9* gene (28). Further, treatment with exogenous BMP ligands was beneficial as it inhibited the pro-inflammatory phenotype of RA FLS (26, 29). Here, we identified that *BMP4* and *SMAD9* are a part of the long-term inflammatory memory that downregulates their expression. Taken together, we speculate that the sustained repression of BMP signaling may be critically required to ensure the persistently transformed phenotype of the RA FLS.

Another interesting result in our study is the identification of *ID4* gene as an SRG. ID4 belongs to the family of the inhibitor of DNA binding domain (ID) containing transcriptional regulators with pivotal roles during development, adult tissue homeostasis and tumorigenesis (30). However, their roles in FLS and RA progression are not known. ID proteins are targets of the BMP signaling (31), and BMP-SMAD-ID signaling network was previously reported to play a role in tumor growth and angiogenesis (32-34). Interestingly, the BMP-SMAD-ID axis was reported to suppress p16/INK4A-mediated cell senescence during the reprogramming of fibroblasts into pluripotent stem cells (35). Since cellular senescence is known to contribute to the increased release of catabolic and inflammation promoting molecules by several cell-types, including the FLS (36), our data suggest that the sustained repression of the BMP4-SMAD8-ID4 axis may function as another unique means to bolster the catabolic phenotype of the RA FLS.

In summary, this study shows that multiple signaling networks are irreversibly modified due to the TNF-mediated long-term epigenetic and transcriptomic reprogramming of the genes in these networks. These genetic factors may together contribute to the persistence in FLS transformation and inflammatory joint diseases. Identification of similar mechanisms in mouse and human FLS suggest that mouse models serve as valuable tools to study at least some of the aspects of therapeutic resistance in inflammatory joint diseases. Future studies such as genetic manipulation of the identified SAGs and SRGs in mice will help establish their pathophysiological significance in RA. Therapeutic targeting of the signaling pathways identified in this study could provide a safer alternative to the use of epigenetics modifying drugs, which are often associated with off-target side effects.

## Methods

### Mouse FLS cultures

FLS were prepared from wild type C57BL/6 mice. The institutional animal care and use committee (IACUC) at Emory University approved animal husbandry and experimental procedures. FLS were isolated by digesting interphalangeal joint synovium according to previously described protocol (11). Hind paws from wild-type mice with a C57/B6 background were deskinned, followed by digestion with 1mg/ml collagenase IV (Sigma-Aldrich) for 2h at 37°C under sterile conditions (35). Cells were cultured in DMEM with 10% fetal bovine serum (Invitrogen) and utilized at passage-4.

### RNA-seq and data analysis

Mice fibroblast cells (FLS) treated with 5 ng/mL recombinant human TNF (R&D Systems) for 16 hours were harvested at 95% confluence. Total RNA was extracted and purified using the RNeasy Mini Kit (Qiagen). Only samples with an RNA integrity number (RIN) >7.5 were used. Libraries were generated from 250 ng RNA using TruSeq Stranded Total RNA Sample Prep Kit (Illumina). Sequencing was carried out using Illumina Hi-Seq 2500 System (Genomics Core Facility, University of Chicago). Except for mouse FLS treatment, all other RNA-seq data were downloaded as FastQ files (**Table S1**) and analyzed using Strand NGS RNA-seq pipeline. Single-end reads were mapped to the m10 mouse genome assembly. Ensembl tools were used for transcript annotation and gene expression analysis. RNA levels were normalized using reads per kilobase of exon model per million mapped (RPKM) sequence method. RNAs whose level was ≥3 normalized RPKM were considered significantly present. For analyses of differential expression levels between mice FLS treated with and without TNF, the cut-off was set ≥ 2 fold change with a *p*-value <0.05 (Audic Claverie test) followed by Benjamin-Hochberg multiple testing corrections for determining false discovery rate.

### ATAC-seq and DNase-seq data analysis

About 50,000 mFLS at passage 4 were treated with or without recombinant human TNF (5ng/mL) for 16h and were processed according to standard ATAC-seq protocol described previously (37). Briefly, cells were lysed, followed by transposition with Tn transposase (Illumina) by PCR amplification using Illumina sequencing adaptor primers (Illumina). Paired-end sequencing was performed on a Hi-Seq 2500 System (University of Georgia Genomics Core) to generate data in FastQ file format. In order to study chromatin accessibility changes due to chronic inflammation, we downloaded FastQ files from a DNase I hypersensitivity sequencing (DNase-seq) experiment of wild-type FLS and inflamed FLS from inflammatory arthritis mouse model (**Table S1**). FastQ files from the ATAC-seq and DNase-seq experiments were analyzed using the Strand NGS sequence analysis pipeline first by aligning the sequences to the mouse genome, mm10 build, followed by removal of duplicate reads. Peak calling and gene annotation were performed by MACS2 algorithm using a padding distance of 5 kb from the transcription start sites of genes.

### Pathway analysis

Ingenuity Pathway Analysis tool was used to predict the canonical regulatory pathways and diseases and functions associated with the SAGs and SRGs (38). Functional protein-protein analysis was performed using STRING protein-protein interaction plugin on Cytoscape, an open source software platform for visualizing complex networks pathway analysis tool (39, 40). Open-source tool, ChEA3 that uses ChIP-seq experiments from the ENCODE database was used to predict transcription factors associations with SAG and SRGs (17).

### Data availability

Accession numbers of the data downloaded from the NCBI-Sequence Read Archive (SRA)-database are indicated in Table S1. RNA-seq and ATAC-seq data generated for this study will be submitted to the NCBI-SRA database.

## Acknowledgements

We thank the support from the National Institutes of Health/National Institute of Arthritis, Musculoskeletal and Skin Disease grant (AR070736) and Startup funds from the Department of Orthopaedics, Emory University School of Medicine to PB.

## Author information

### Affiliations

Department of Orthopaedics and Department of Cell Biology, Emory University School of Medicine, Atlanta, GA 30322, USA: Umesh Gangishetti, Sergio Ramirez-Perez, Kyle Jones, Abul Arif, Hicham Drissi and Pallavi Bhattaram.

Atlanta VA medical Center, 1670 Clairmont Rd, Decatur, GA 30033: Hicham Drissi.

### Contributions

PB conceived the study, performed experiments, analyzed data contributed to manuscript writing; HD contributed to manuscript writing; UG performed experiments; SR contributed to conceptual design and manuscript writing. AA contributed to data analysis, preparation of figures and manuscript writing.

Correspondence and requests for materials should be addressed to P.B.

## Additional information

The manuscript contains 6 Figures, 1 Supplementary Figure and 4 Supplementary tables.

