## Supplementary material for "Chronic exposure to TNF reprograms cell signaling pathways in fibroblast-like synoviocytes by establishing long-term inflammatory memory": Fig S1

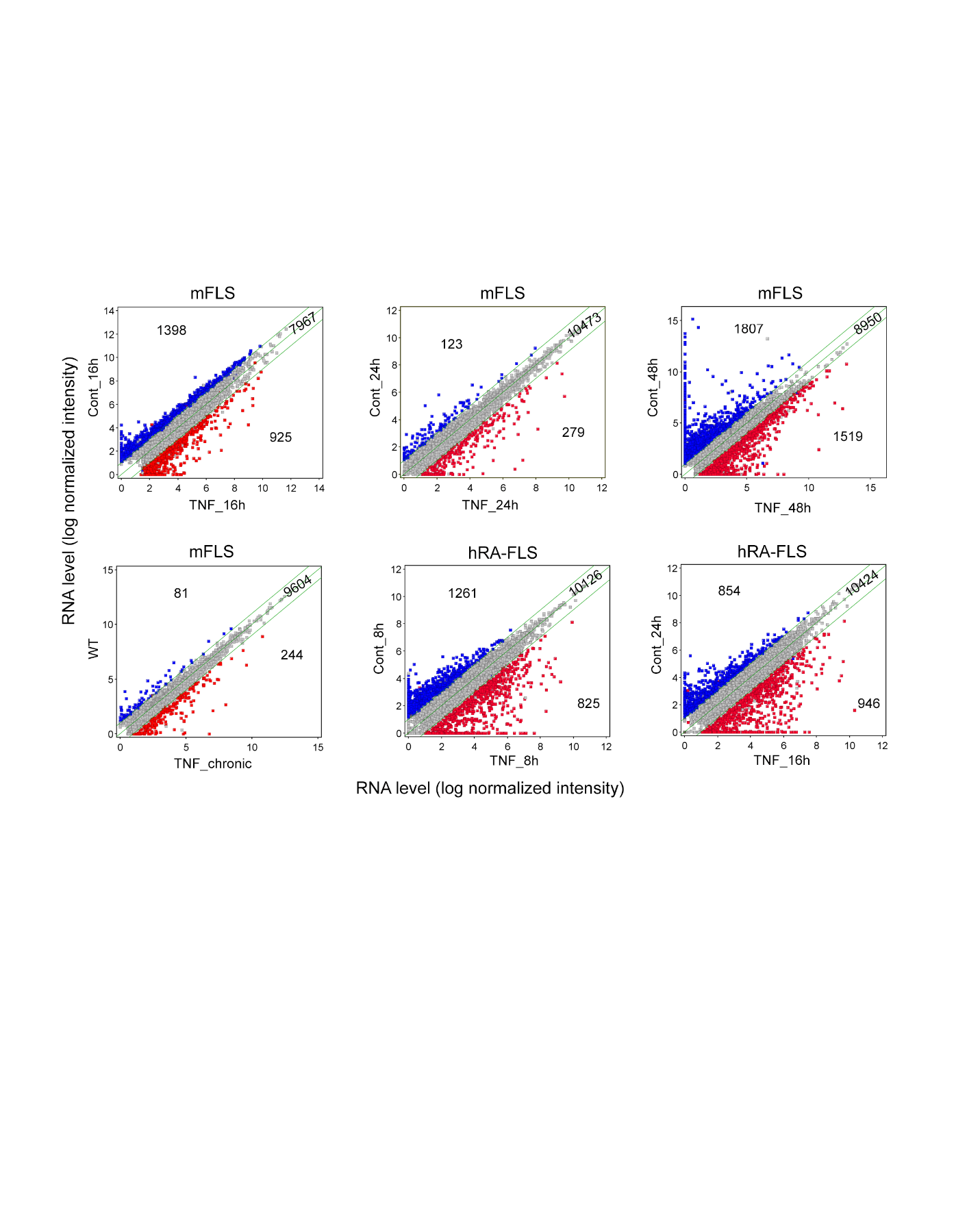

**Supplementary Figure 1.** Scatter plots of RNA-seq data. Averaged normalized intensities from various experimental groups are presented. Green lines indicate 2-fold chage.
